# Differential transcriptomics and metabolomics analyses of the skin of coarse and fine Liaoning cashmere goats

**DOI:** 10.1101/2025.08.13.670025

**Authors:** Jianwei Wei, Zuo Shi, Na Liu, Di Han, Chunqiang Wang, Wei Ma

**Affiliations:** College of Animal Husbandry and Veterinary Medicine, Jinzhou Medical University, Liaoning 121001, China; Liaoning Modern Agricultural Production Base Construction Engineering Center, Ministry of Animal Husbandry, Liaoning 111000, China

## Abstract

Secondary follicles in cashmere goats determine cashmere yield and quality. The present study investigated the relationship between the regulatory factors and cashmere fiber diameter through transcriptomics and metabolomics methods. Diameters of the fibers of 400 (18 months old) female Liaoning cashmere goats were measured and categorized into coarse and fine hair types (200 goats per category). Six goats were selected as representatives of each group for further analysis. Fiber diameters were significantly different between groups (*P* < 0.05). Transcriptomic analysis revealed 211 differentially expressed genes between groups, including *FGF18*, *KRT36*, *KRT79*, *AWAT2*, and *MOGAT1*, which were enriched in the PPAR and glycerol–lipid metabolism signaling pathways that regulate hair follicle growth and sebum secretion. Metabolomics analysis revealed 50 differentially expressed metabolites in the skin, which regulate cellular ion channels and participate in neurological channels by enriching ABC transporter protein pathways, amino acid biosynthesis and metabolism, and arachidonic acid metabolism. The findings of the present study could facilitate improvement of cashmere quality in cashmere goats.

## Introduction

Liaoning cashmere goats are associated with high cashmere production, high net cashmere content, and stable performance after selective breeding. Cashmere goat skin contains two types of hair follicles: primary hair follicles (PFs) for coarse hair growth and secondary follicles (SFs) for villus or fine hair growth [1]. The growth and development of SFs influence cashmere yield and quality, which are regulated by various genes and signaling and metabolic pathways. Certain metabolites are crucial in hair follicle development and cashmere quality [2]. Histological research techniques are widely used to analyze the key genes and metabolites associated with cashmere quality, which could promote high-quality cashmere production [3]. Transcriptomics, based on high-throughput sequencing technologies, can be used to comprehensively analyze the gene expression profiles of specific tissues or cells during particular periods. It has been applied extensively in research such as the cashmere growth cycle and hair follicle development in cashmere goats. For instance, Zhang et al. [4] employed RNA-seq technology to identify key differentially expressed genes (DEGs) during the growth of SFs in cashmere goats, providing crucial insights into the regulatory mechanisms of hair follicle development. Metabolomics, conversely, detect the types and quantities of all small molecule metabolites within an organism, reflecting physiological status and metabolic pathways. In addition, Li et al. used Liquid Chromatography-Tandem Mass Spectrometry to analyze metabolite changes in skin tissues of cashmere goats at different growth stages, revealing key metabolic pathways associated with cashmere growth [5].

However, single-omics analysis cannot unravel the regulatory networks of complex biological processes adequately or comprehensively. The integration of transcriptomics and metabolomics can combine gene expression and metabolite variation information to facilitate construction of gene-metabolite regulatory networks, offering greater insights into the molecular mechanisms underlying biological processes.

Hair fineness in cashmere goats is a key factor influencing the quality of cashmere. After prolonged selective breeding, cashmere yield and quality in Liaoning cashmere goats have improved; however, there is potential for further improvement. As hair follicle development in skin tissue influences both cashmere fineness and yield, identifying the regulatory factors and genes involved in hair follicle growth and development has become a major research focus.

Therefore, the aim of the present study was to systematically identify key genes and metabolic pathways influencing cashmere yield and quality in cashmere goats using integrated transcriptomic and metabolomic analysis. The findings could provide a theoretical basis for molecular breeding and nutritional regulation, thereby promoting the sustainable development of the cashmere goat industry.

## Materials and methods

### Animals and experimental design

Eighteen-month-old Liaoning cashmere goats were obtained from two non-inbred lines. The goats had the same rearing conditions and similar body weights. They were all obtained from the same field (Liaoning Modern Agricultural Production Base Construction Center, Liaoyang, China). The Animal Ethics Committee of Jinzhou Medical University (Jinzhou, China) approved all methods and procedures involved in this experiment (Authority no.: 20240523).

A total of 400 female Liaoning cashmere goats were classified into coarse and fine groups based on cashmere diameter, with 200 goats in each group. The classification was based on the Chinese Cashmere Classification Standard GB18267-201: 15.5 μm < cashmere diameter ≤ 16.0 μm was defined as fine, and 16.0 μm < cashmere diameter ≤ 18.5 μm was defined as coarse.

For the collection of fleece samples, a 5 × 5 cm^2^ area was selected from the right scapula of the goat, and the fleece in the area was cut close to the skin using curved scissors. The fleece was then stored in a sealed bag in its natural state and labeled to determine the diameter of the cashmere fibers.

For skin collection, samples were collected along the upper edge of the scapula using a biopsy perforator with a diameter of 1 cm. The samples were frozen quickly in liquid nitrogen and stored at −80 °C for later use.

The cashmere samples were tested, cleaned, and processed after being dried naturally. An N-107CCD Wool Fineness Tester (Ningbo Yongxin Optical Co., Ltd., Ningbo, China) was used to determine the diameters of the cashmere fibers. Based on the extreme values, six goats per group with comparable baseline traits were selected for follow-up histological analysis.

### Transcriptomics analysis of skin tissue

Total RNA was extracted from the 12 tissue samples. Agarose gel electrophoresis was used to determine RNA integrity. The RNA concentration and purity were determined using a Nanodrop 2000 spectrophotometer (Thermo Fisher Scientific, Waltham, MA, USA). RNA concentration was quantified accurately using a Qubit 2.0 Fluorometer (Thermo Fisher Scientific). The RNA integrity number value was determined with an Agilent 2100 Bioanalyzer (Agilent Technologies, Santa Clara, CA, USA) for precise determination of RNA integrity. The RNA-seq library was constructed using the Illumina TruSeq^TM^ RNA Sample Prep Kit (Illumina, San Diego, CA, USA) and sequenced with the Illumina high-throughput sequencing platform (Illumina). FastP software was used to perform quality control and evaluation of raw data. Using Hisat2 [6,7], the filtered reads were compared with the reference genome (version: GCF_001704415.1 https://ftp.ncbi.nlm.nih.gov/genomes/refseq/vertebrate_mammalian/Capra_hircus/). EdgeR was used to identify genes with differential expression between samples. Genes with a false discovery rate (FDR) ≤ 0.05 and a |log2 FC| ≥ 1 were considered DEG candidates. ClusterProfiler was employed for enrichment analysis of the functions and pathways of the DEGs. The sequences were imported into the GO database for comparison and classification (http://www.geneontology.org/) and then imported into the KEGG database (http://www.genome.jp/kegg/) to determine the biological functions of the genes at the system level.

### Metabolomics analysis of skin tissue

The 12 skin samples of 25 mg each were placed in Eppendorf tubes (Eppendorf, Hamburg, Germany). Subsequently, extraction, vortexing, grinding with a steel ball, and ultrasonication in an ice-water bath were performed (four replicates). After centrifugation, the supernatants were transferred to Eppendorf tubes, and a small amount of each sample was mixed to establish a quality control sample. The extract was dried and processed. Subsequently, a methoxylamine salt reagent was added, and the mixture was mixed and incubated in an oven. Bis (trimethylsilyl)trifluoroacetamide was then added, and the mixture was incubated at a high temperature (70 °C) for 1.5 h. After equilibration at room temperature (20–25 °C), fatty acid methyl esters were added to the samples, which were then subjected to an uptake test in a random order. Gas Chromatography-Time-of-Flight-Mass Spectrometry analysis was performed using an Agilent 7890 gas chromatograph (Agilent Technologies), coupled with a time-of-flight mass spectrometer. Raw data analysis, including peak extraction, baseline adjustment, deconvolution, alignment, and integration, was conducted using Chroma TOF v4.3x (LECO Corp., St. Joseph, MI, USA), and the LECO-Fiehn Rtx5 database (https://fiehnlab.ucdavis.edu/projects/fiehnlib) was used for metabolite identification by matching the mass spectra and retention indices. Finally, the peaks detected in less than a half of QC samples or RSD > 30% in QC samples was removed [8].

A *P*-value < 0.05 in the Student’s t-test and a Variable Importance in the Projection (VIP) value > 1 in the first principal component of the Orthogonal Partial Least Squares-Discriminant Analysis model were used as screening criteria for differential metabolites.

### qRT‒PCR verification

To verify the accuracy of the transcriptomic sequencing data, four candidate genes, including *KRT9*, *FGF18*, *SEMA3D,* and *FICD*, were selected randomly for qRT-PCR validation in the six other skin samples from the coarse group (n = 200). Total RNA was extracted, and cDNA was synthesized using a Total RNA Reverse Transcriptase Kit (Takara, Kusatsu, Japan). cDNA was prepared using SYBR Premix Ex Taq™ II (Takara), and quantitative PCR was conducted on an ABI 7500 Fast Real-Time PCR system (Applied Biosystems, Waltham, MA, USA). The optimized cycling conditions were as follows: denaturation at 94 °C for 5 min, 94 °C for 15 s, and 55 °C for 15 s. For each sample, 45 cycles were performed in triplicate. Relative expression was determined using the 2^−ΔΔCT^ method, and the results were normalized using *GAPDH* [9–11] as an internal reference gene. The primer sequences are listed in S1 Table.

### Statistical analysis

Cashmere fiber diameter is expressed as the mean ± standard deviation. IBM SPSS Statistics 25 (IBM Corp., Armonk, NY, USA) was used for comparisons between two samples, and *P* < 0.05 indicated a statistically significant difference. IBM SPSS Statistics 25 software (IBM Corp.) was also used to conduct correlation analysis between DEGs and metabolites.

## Results

### Measurement of cashmere-related indices and selection of individuals for further testing

Fiber diameter was determined and analyzed using cashmere samples from 400 ewes. A significant difference was observed in average fiber diameter between the two groups of cashmere samples (*P* < 0.05) (Table 1).

**Table 1.** Measurement results of cashmere-related characteristics. Data in the same column marked with different lowercase letters indicate a significant difference (*P* < 0.05). No letters indicate a non-significant difference (*P* > 0.05).

| Colony | Sample volume | Fiber diameter<br>($\mu\text{m}$) | Largest<br>diameter ($\mu\text{m}$) | Smallest<br>diameter ($\mu\text{m}$) |
| --- | --- | --- | --- | --- |
| Group I | 200 | $17.57 \pm 0.56^a$ | 19.82 | 16.33 |
| Group II | 200 | $15.42 \pm 0.37^b$ | 16.23 | 14.04 |

Group I: Coarse group
Group II: Fine group

Based on the diameter values of cashmere fibers in the coarse and fine groups, the skin tissues of six goats were randomly selected from each group for subsequent transcriptomic and metabolomic analyses.

### Transcriptomics analysis of skin tissues

#### DEG analysis

After removing adapters and low-quality reads from the raw data, an average of 19610092 bp or more of clean reads was obtained for each sample, such that the minimum proportion of clean reads in the sample was 89.25%. Q20 and Q30 were at least 98.71% and 95.16%, respectively, and the GC content was at least 50.36% (S2 Table), which meets the requirements for further bioinformatics analysis.

Differentially expressed genes in each group were analyzed using EdgeR and corrected using multiple hypothesis testing, log2FC was used as a quantitative measure to evaluate differences in gene expression levels. By comparing coarse to fine types, a total of 211 DEGs were identified, wherein 75 and 136 were significantly upregulated and downregulated, respectively (log_2_FC value > 0 indicates gene upregulation, meaning the expression level was significantly higher in the coarse group than in the fine group. Conversely, a value < 0 indicates downregulation, meaning the expression level was significantly lower than that in the fine group) (Fig 1). The DEG analysis results are presented in Table 2 (Top 10, based on log_2_FC value), and all DEG results are presented in S1 File.

**Fig 1.**
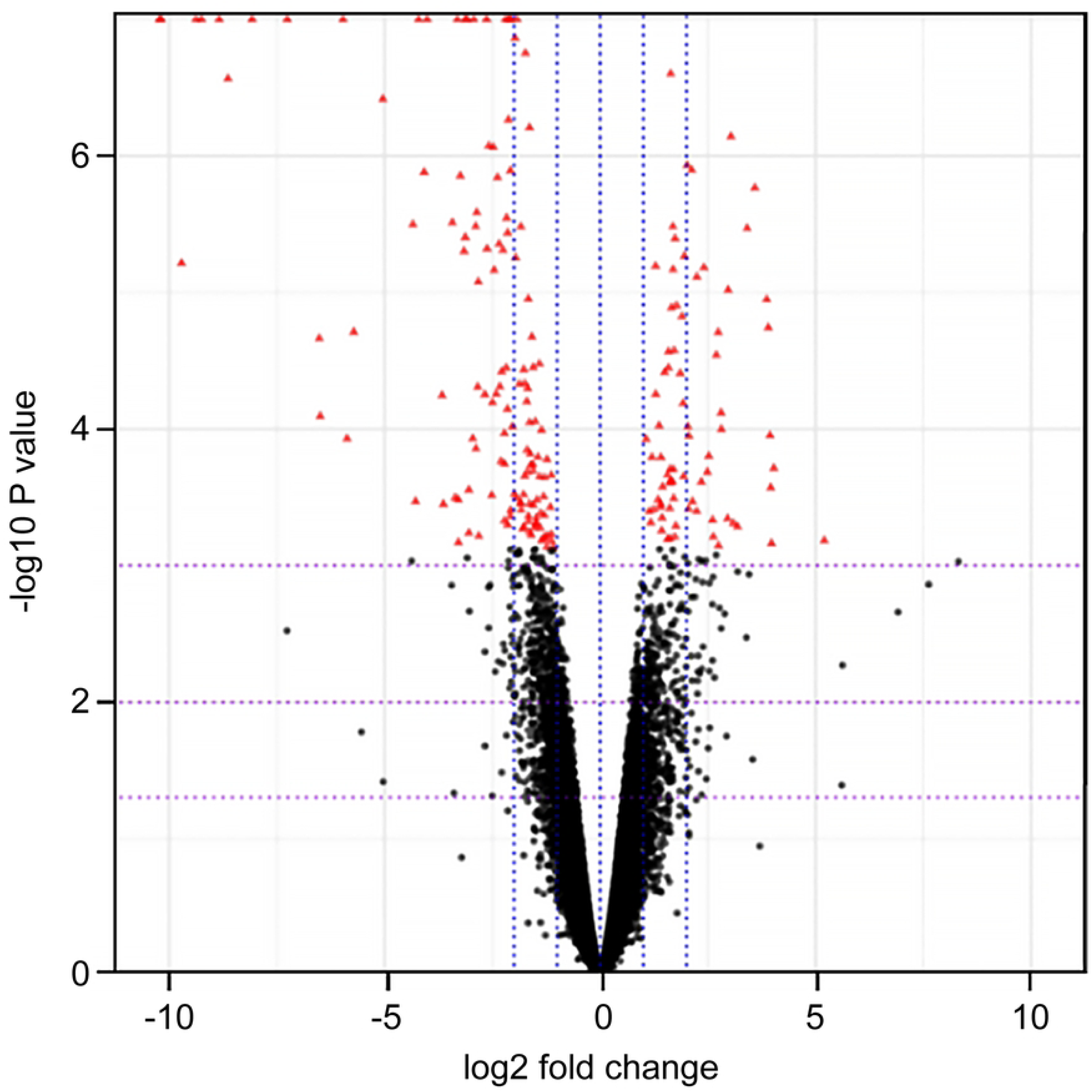
Volcano plot of differentially expressed genes. Each point represents a gene; the horizontal axis indicates the log2 value of the difference in diversity; the vertical axis indicates the negative logarithmic value of the *P*-value; the red dots in the figure represent the genes for which the statistical test showed differences.

**Table 2.** Differentially expressed genes in the skin of coarse and fine goat types (Top 10). Gene_ID: Gene ID; Fpkm_sample: fragments per kilobase of transcript per million mapped reads (Fpkm) value of the sample; log2FC: values > 0 represent upregulation and less than 0 downregulation; *P-*value: the level of significance of the calculated statistical difference test; FDR: the result of the correction of a multiple hypothesis test for *P-*value.

| Gene ID | Fpkm | Fpkm | log<sub>2</sub>FC | *P*-value | FDR | Expression |
| --- | --- | --- | --- | --- | --- | --- |
|  | ( I ) | ( II ) |  |  |  | level |
| *UCN3* | 25.32 | 3.77 | 3.03 | 7.21E-07 | 0.00036419 | up |
| *AMH* | 1.91 | 0.28 | 2.97 | 9.59E-06 | 0.00218683 | up |
| *SOAT1* | 5.07 | 114.03 | -4.20 | 6.65E-12 | 9.41264698 | down |
| *ADAMTS4* | 0.44 | 5.31 | -3.30 | 8.80E-11 | 1.13194921 | down |
| *CES3* | 2.66 | 19.37 | -2.58 | 8.44E-07 | 0.00040568 | down |
| *LNPEP* | 0.69 | 4.62 | -2.46 | 6.86E-06 | 0.00164492 | down |
| *LCLAT1* | 1.47 | 8.41 | -2.14 | 7.16E-05 | 0.01107718 | down |
| *TGOLN2* | 5.79 | 29.18 | -2.03 | 9.61E-05 | 0.01321013 | down |
| *ATP11C* | 1.96 | 7.50 | -1.63 | 8.94E-05 | 0.01264619 | down |
| *ITPR2* | 1.50 | 5.14 | -1.50 | 8.79E-05 | 0.01256360 | down |

#### Gene Ontology enrichment analysis

Gene Ontology (GO) functions are classified into three main groups: biological process (BP), cellular component (CC), and molecular function (MF). The enrichment results of upregulated and downregulated genes are shown in Table 3 (a few examples), and all enrichment results are shown in S2 File (level 2, the second level in the GO classification).

**Table 3.**
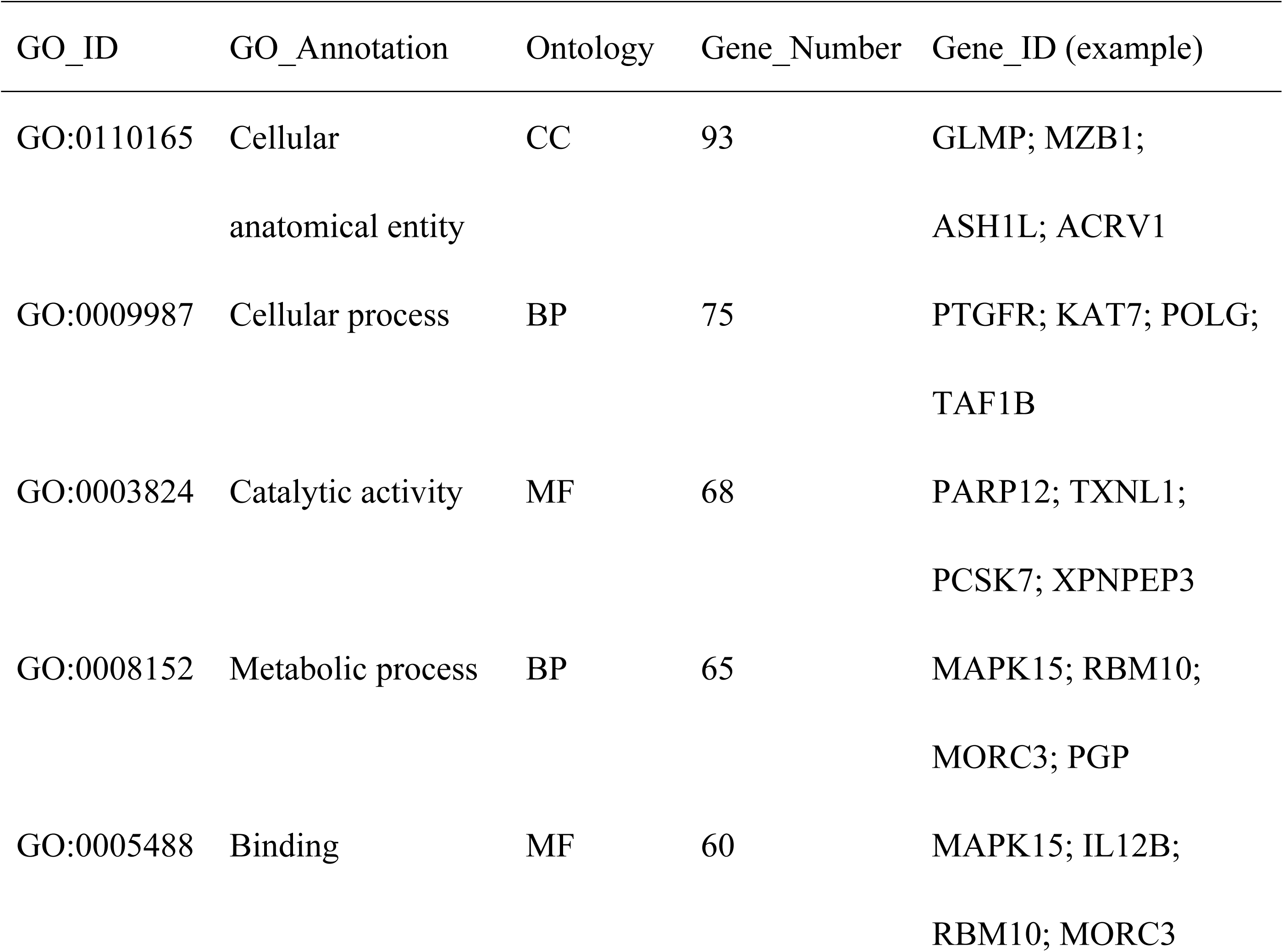
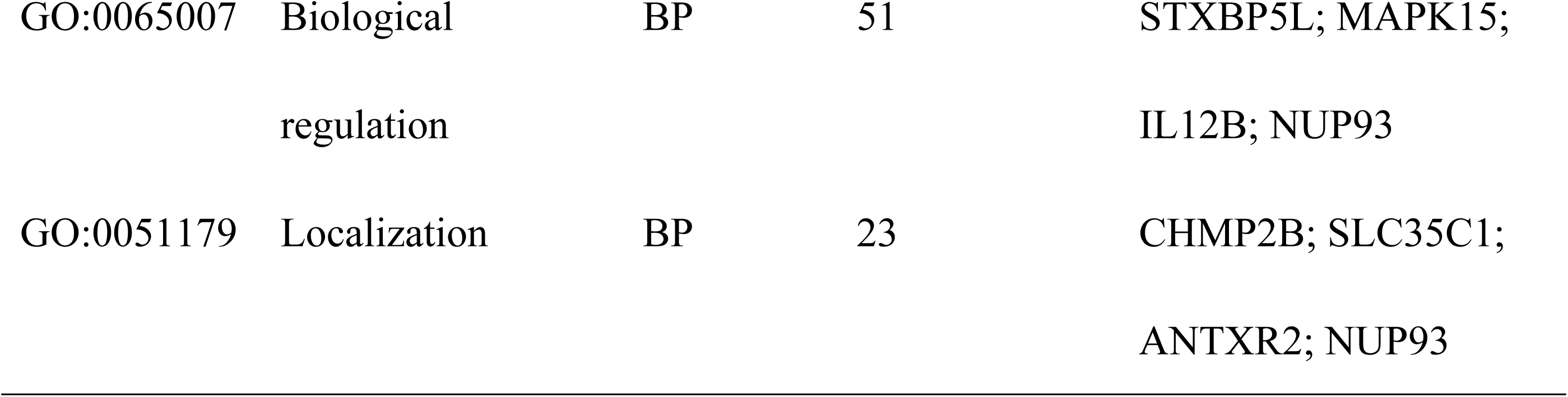
Gene Ontology functional annotation results (Level 2, a few examples). GO_ID: GO Function ID; GO_Annotation: GO Function Description; Ontology: Ontology classification; Gene_Number: Number of genes; Gene_ID: Gene name.

The upregulated DEGs were enriched for 110 BP, 43 CC, and 58 MF terms. In the BP category, genes involved in lipid metabolism, such as lipid synthesis and catabolism, fatty acid elongation, and saturated and unsaturated fatty acid metabolism, showed significant enrichment, followed by nucleic acid metabolism and coenzyme metabolism. In the CC category, several genes showed enrichment, although not significantly, in the endoplasmic reticulum, mitochondria, and lipid droplets.

Downregulated DEGs were enriched for 314 BPs, 105 CCs, and 151 MFs. Among the enriched BPs, redox processes and small-molecule metabolism processes showed relatively higher enrichment levels. Cellular component category enrichment was mainly in the cytoskeleton, rough endoplasmic reticulum, and nuclear transcription factor complex. The major MFs enriched were related to ion metabolism, including iron and chlorine ion metabolism (Figs 2 and 3).

**Fig 2.**
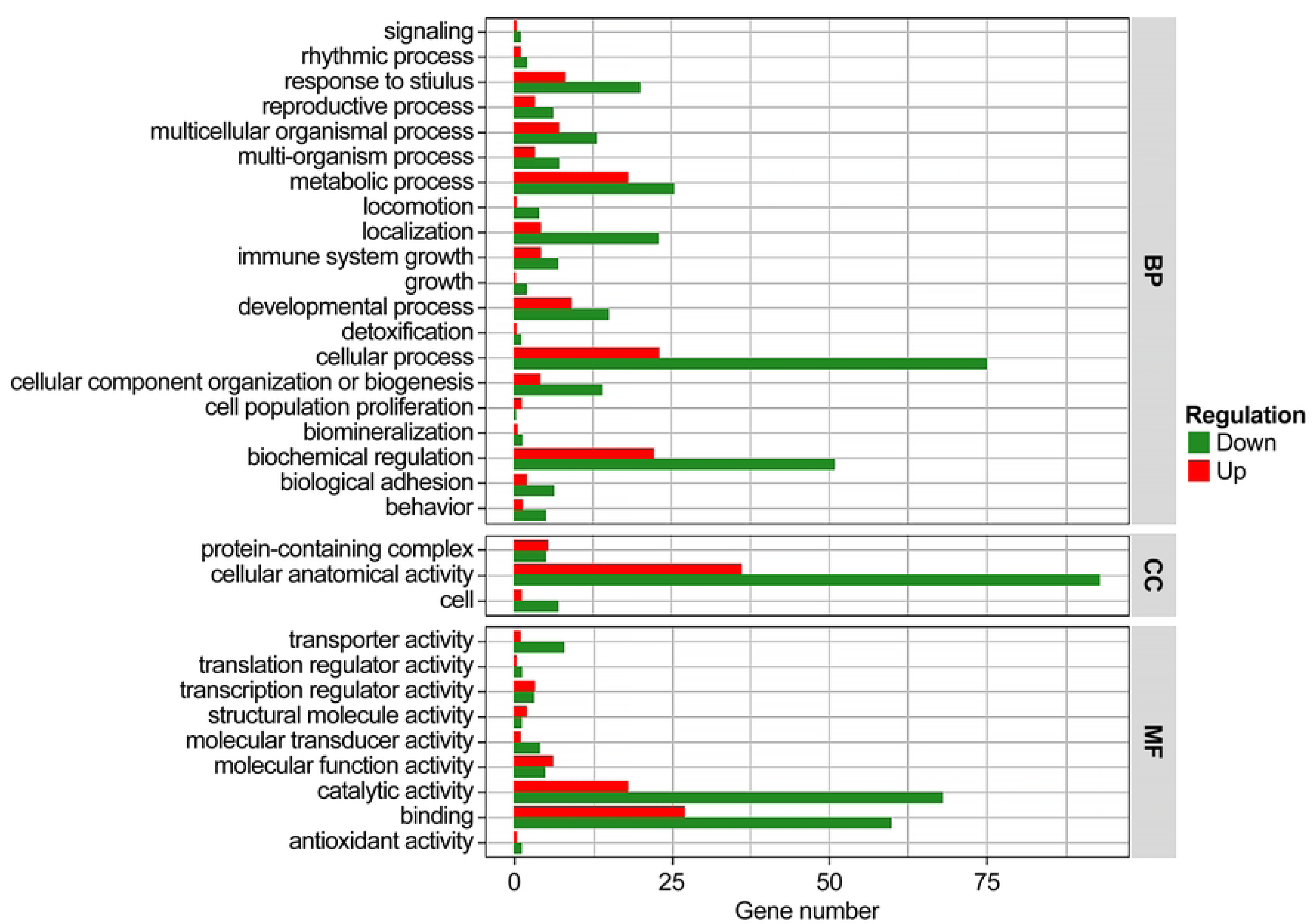
Differentially expressed gene statistics based on Gene Ontology term (Level 2). Numbers of differentially upregulated and downregulated genes in each component of Gene Ontology Term Level 2.

**Fig 3.**
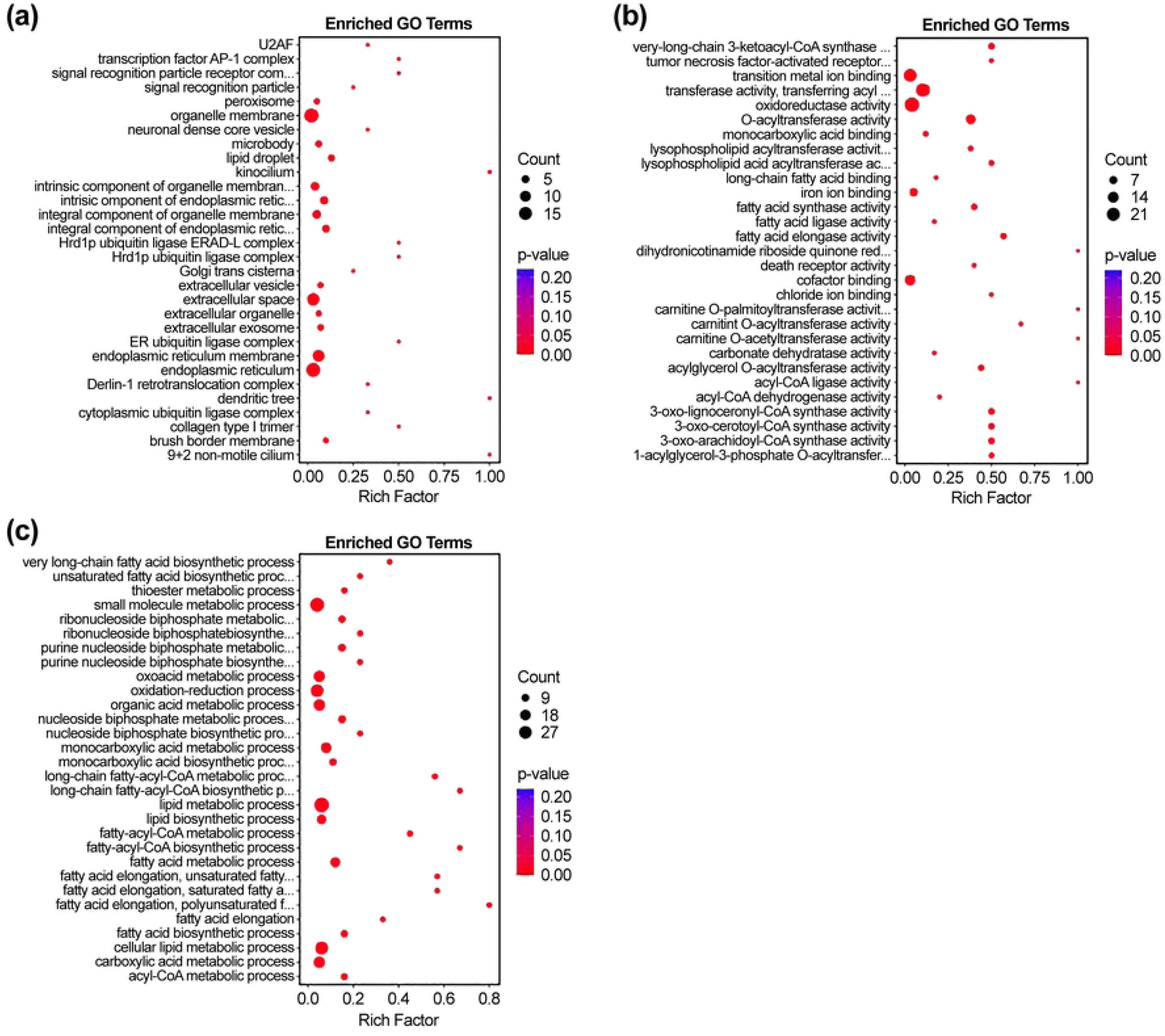
Gene Ontology (GO) analysis of differentially expressed genes (DEGs) (Level 2). Functions of DEGs under different GO enrichment categories: biological processes (a), cellular components (b), and molecular functions (c).

#### KEGG enrichment analysis of DEGs

To further identify the major metabolic and signaling pathways of the DEGs, Kyoto Encyclopedia of Genes and Genomes (KEGG) enrichment analysis was performed. Pathways were considered significantly enriched at *P-*values < 0.05. The DEGS mapped to 245 KEGG pathways, 40 of which were significantly enriched, including PPAR signaling, glycerol–lipid metabolism, retinol metabolism, unsaturated fatty acid biosynthesis, relaxin signaling, adipocytokine signaling, and fatty acid degradation. Among the genes, *MOGAT1* and *AWAT2*, which are involved in glycerol–lipid metabolism and retinol metabolism, respectively, were downregulated significantly (Fig 4).

**Fig 4.**
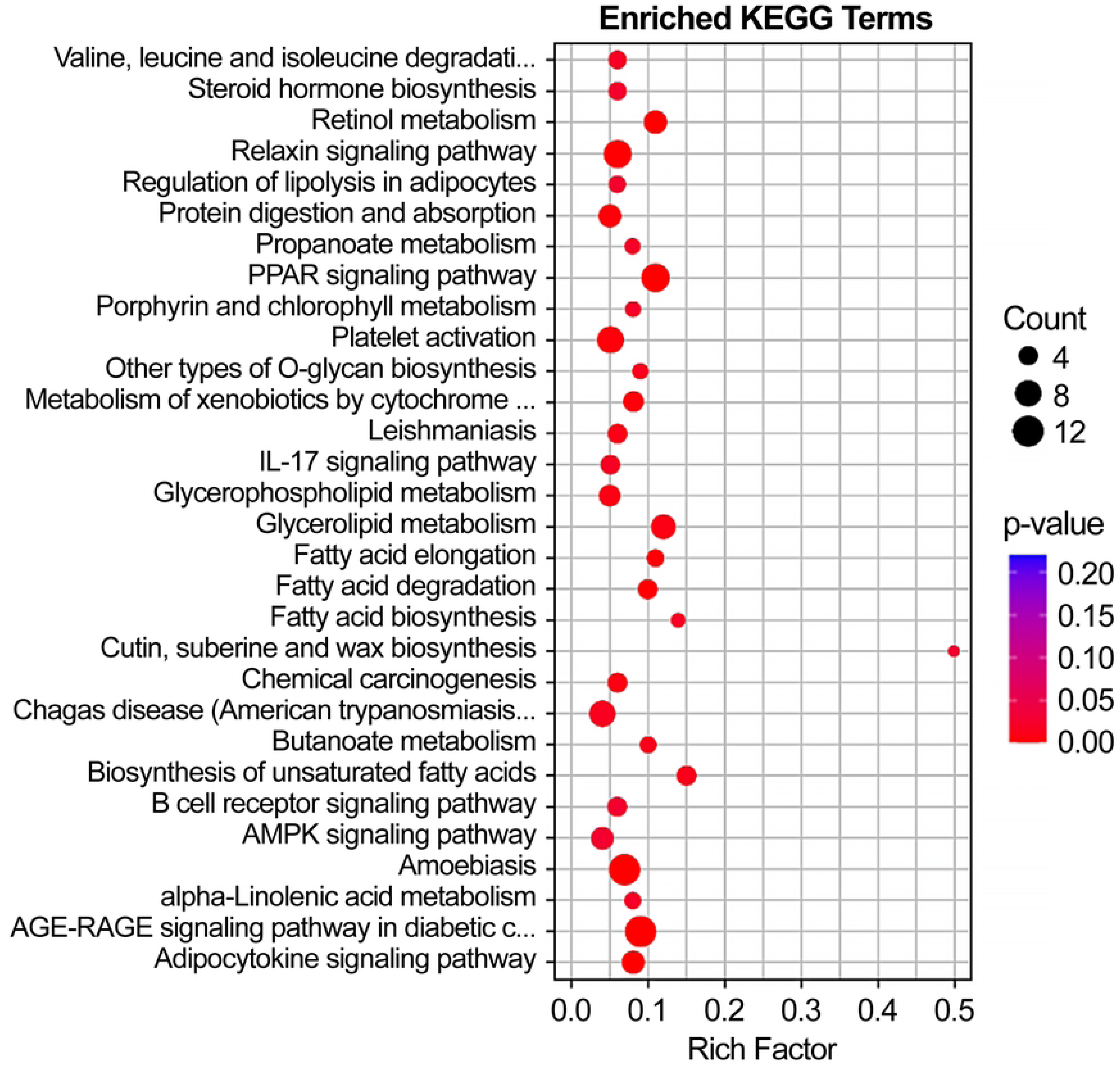
Display of differentially expressed genes (DEGs) in Kyoto Encyclopedia of Genes and Genomes (KEGG) pathways. The red border indicates the presence of upregulated DEGs, whereas the blue border indicates the presence of downregulated DEGs.

#### qRT‒PCR validation of four candidate genes

Four candidate genes were randomly selected for qRT-PCR analysis, including two upregulated genes, *KRT9* and *FGF18*, and two downregulated genes, *SEMA3D* and *FICD*, from other six individuals in the coarse group (n = 200). The results showed that the qRT-PCR expression levels of the four genes were consistent with the RNA-seq results (Fig 5), indicating reliability of the RNA-seq results.

**Fig 5.**
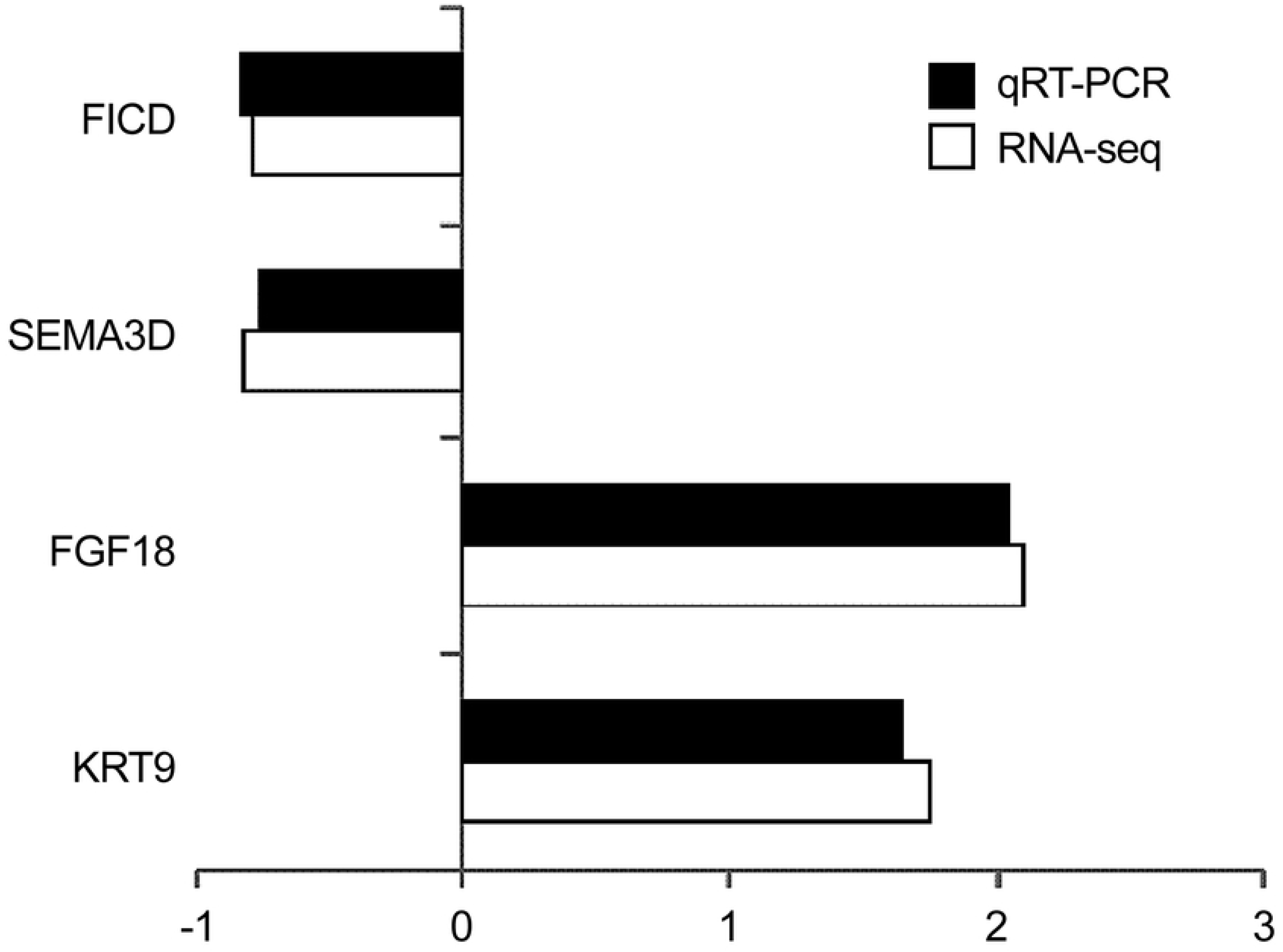
**Comparison of qRT-PCR assay and RNA-seq results for four genes, including *KRT9.*** Black bars represent qRT-PCR detection results, whereas the white bar graphs represent RNA-seq results. Points in the left of the 0.0 value on the horizontal axis indicate downregulation, whereas those to the right indicate upregulation.

### Metabolomics analysis of skin tissues

#### Screening for differentially abundant metabolites

The screening criteria for differential metabolites are VIP > 1 and *P*-value < 0.05. In total, 448 metabolites were detected in the skin tissue of coarse- and fine-type cashmere goats, including 110 differentially abundant metabolites, where 97 and 13 were upregulated and downregulated, respectively (upregulation refers to metabolites with significantly higher abundance in the coarse group compared to in the fine group, whereas downregulation indicates metabolites with significantly lower abundance in the fine group compared to in the coarse group) (Fig 6a). There were 50 metabolites with different abundances, wherein 45 were upregulated, including proline, valine, alanine, isoleucine, oxyproline, glutamate, inosine, L-allothreonine, lactate, serine, and myo-inositol, and five were downregulated, including glucose, mannose, phthalic acid, 3-aminoisobutyric acid, and glucosamine. The screening results are shown in Table 4, and the differences in metabolite expression models of the different samples are shown in Fig 6b, and all differential metabolites are presented in S3 File.

**Fig 6.**
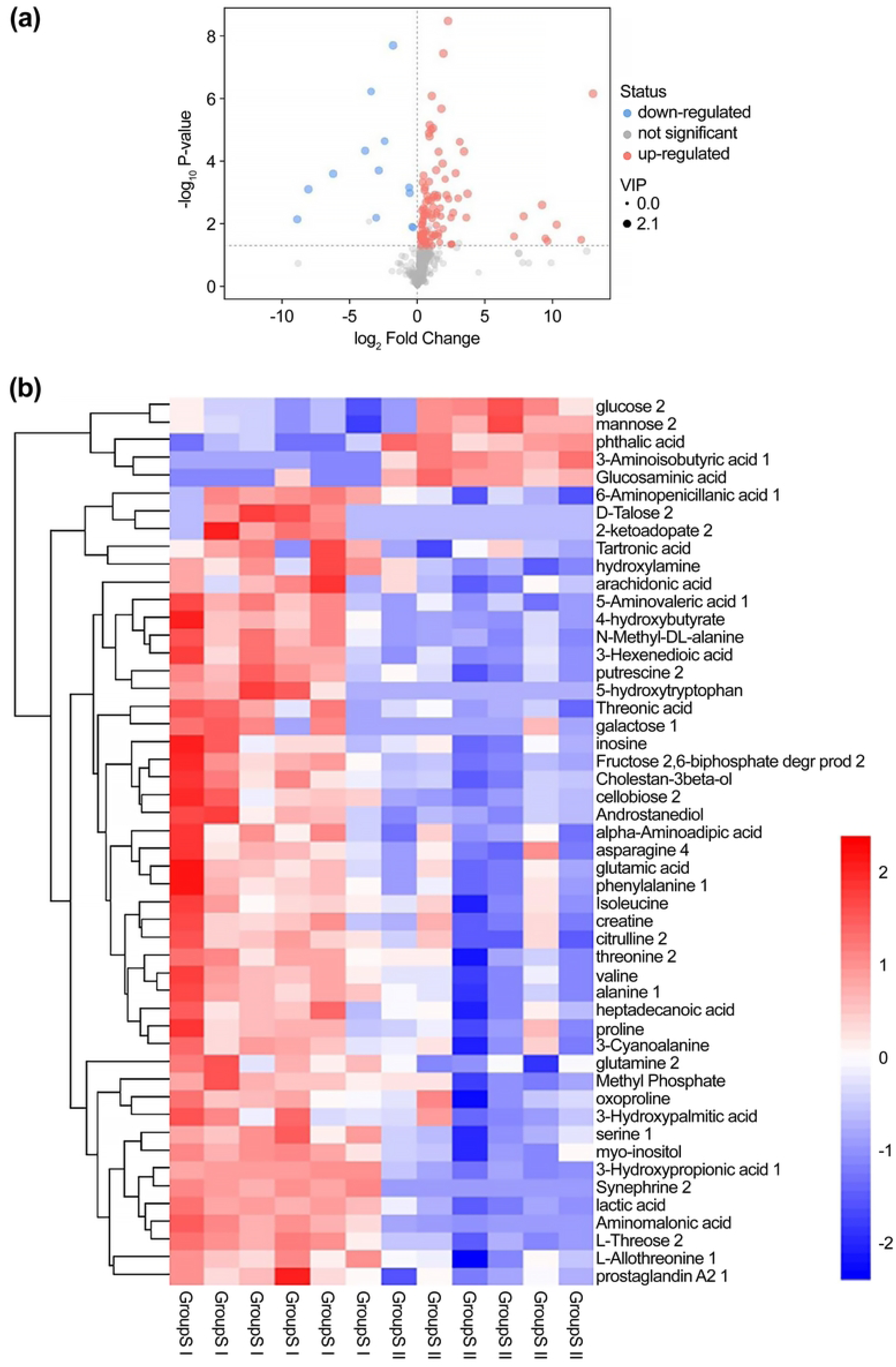
Differentially abundant metabolites in the skin tissue of coarse and fine haired goats. (a) Volcano diagram of all differentially expressed metabolites in the coarse and fine group tissues. (b) Heat map of accumulation of known differentially expressed metabolites in the coarse and fine group tissues.

**Table 4.**
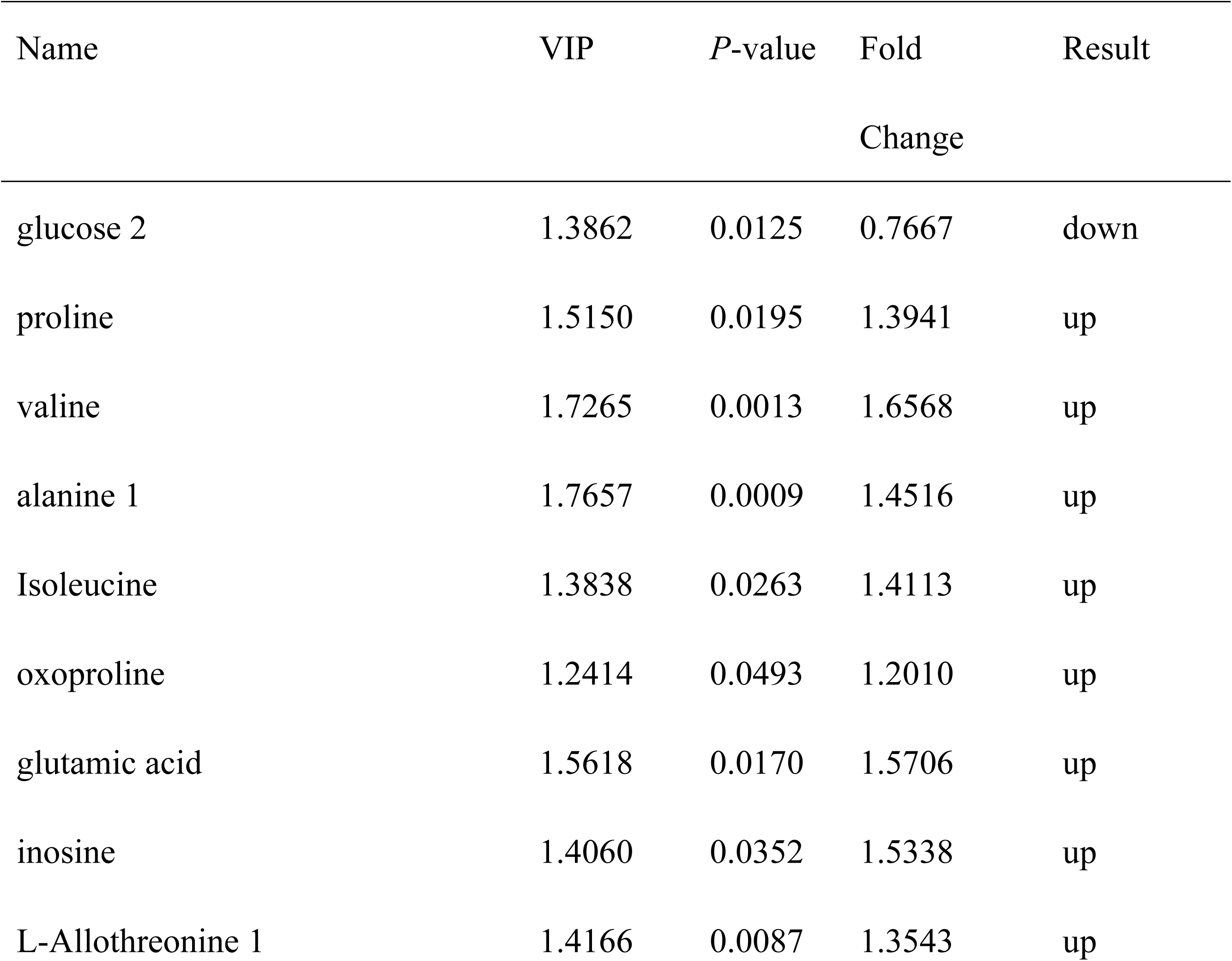
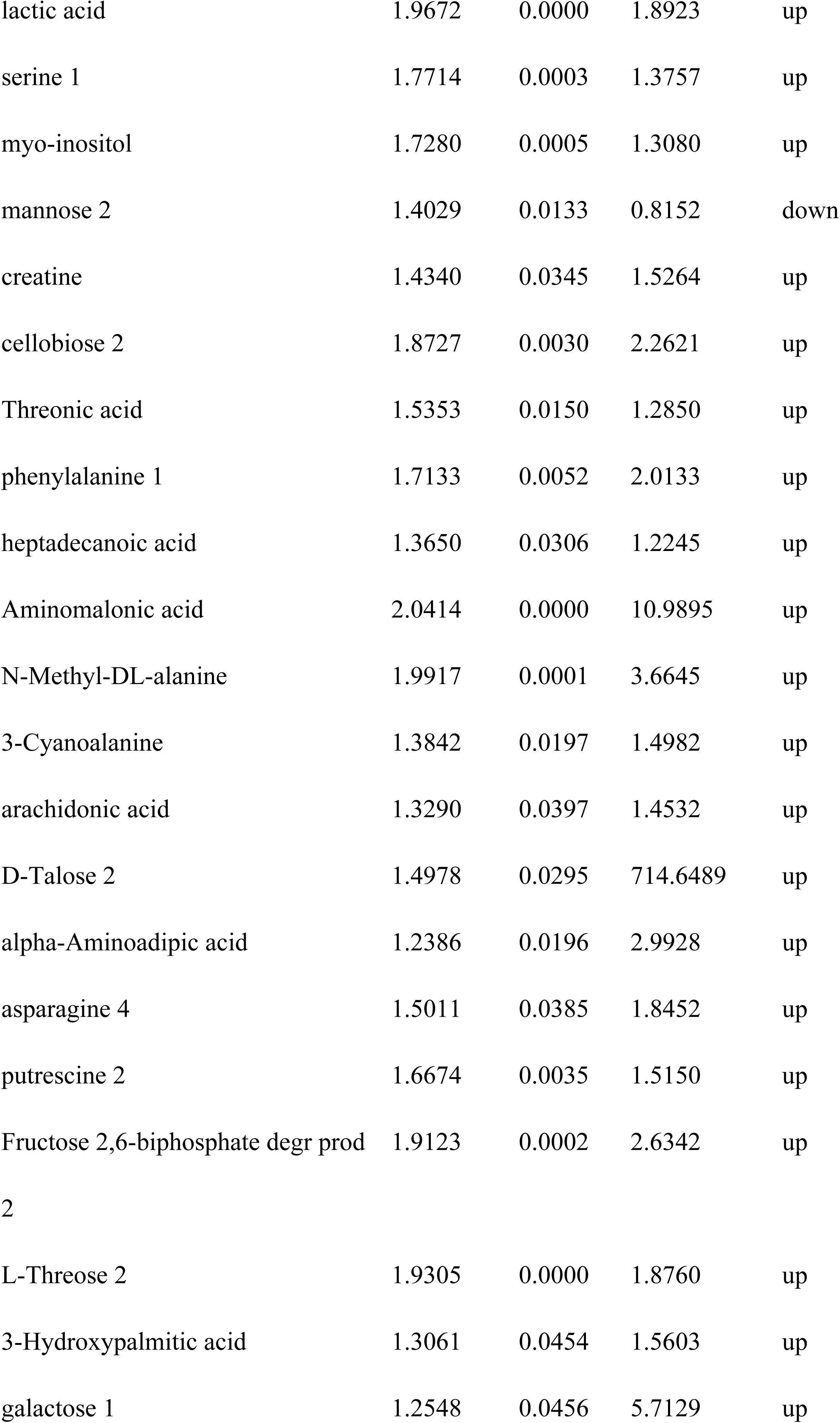
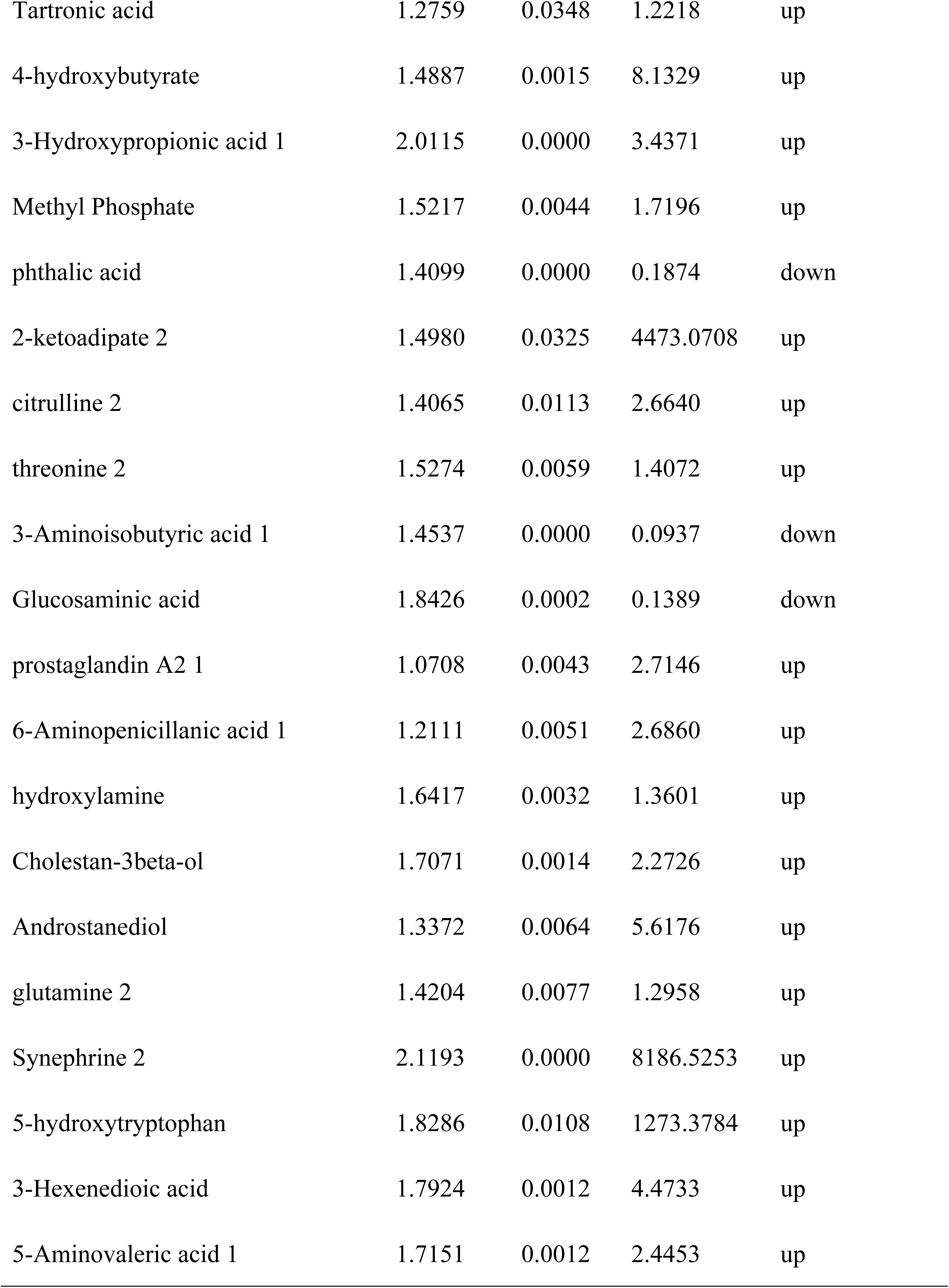
Potential biomarkers affecting cashmere thickness in the skin tissue. Name: Name of the substance in the Fiehn database; VIP: Variable Importance in Projection; *P*-value: *P*-value from the t-test; Fold Change: Quantitative ratio of substances between two groups; Result: Screening results.

#### KEGG pathway analysis of differentially abundant metabolites

In addition to the glycine, serine, and threonine metabolism pathways, skin tissue also hosted ABC transporter, amino acid biosynthesis, and metabolic pathways, as well as other enrichment pathways, including valine, leucine, and isoleucine biosynthesis; arachidonic acid metabolism; mineral intake; protein digestion and absorption; arginine and proline metabolism, leucine and isoleucine metabolism, arachidonic acid metabolism, mineral absorption and protein digestion and absorption. The differentially abundant metabolites that matched some metabolic pathways accurately are listed in Table 5; L-proline and L-valine appeared in several metabolic pathways.

**Table 5.**
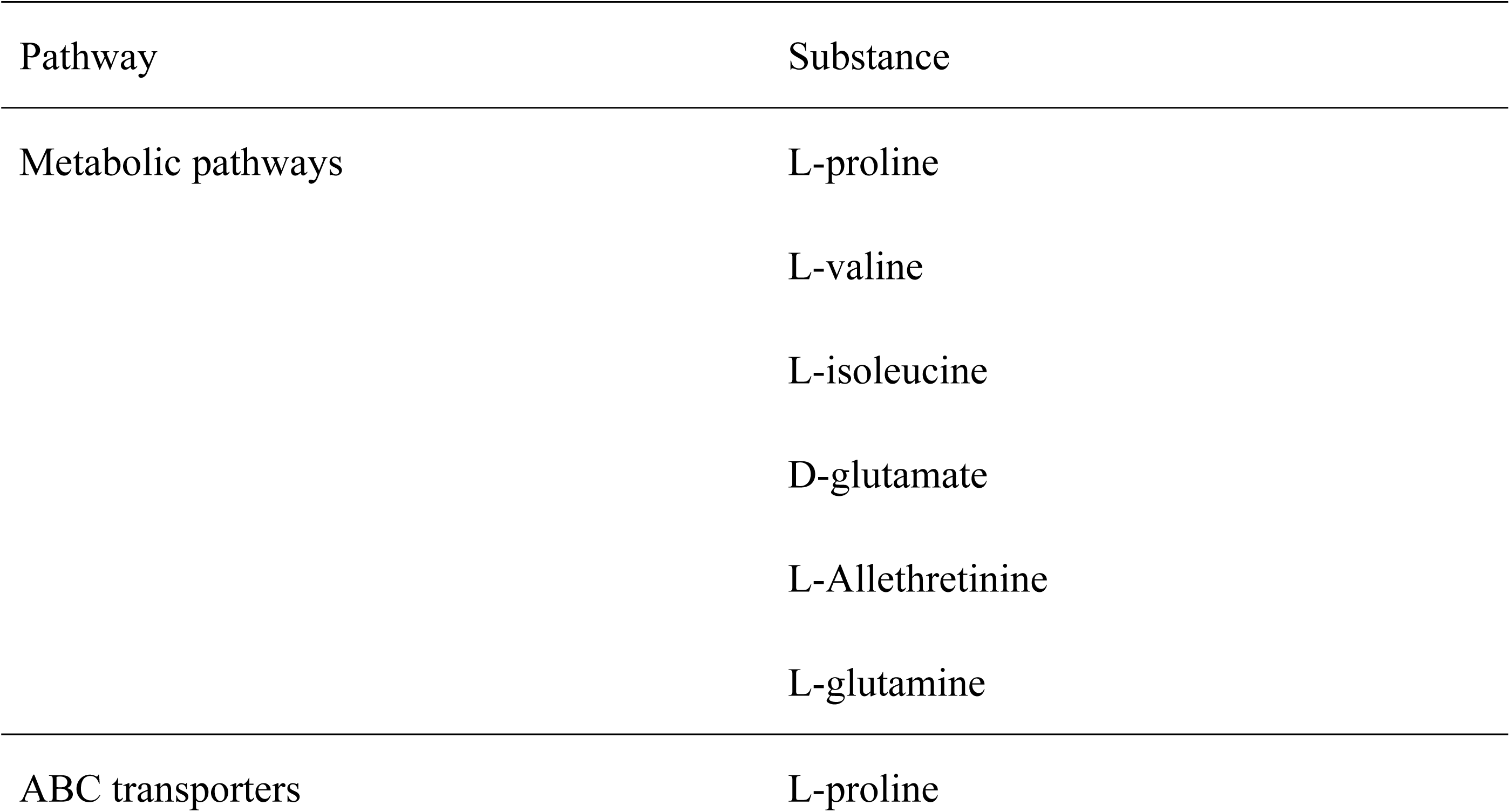
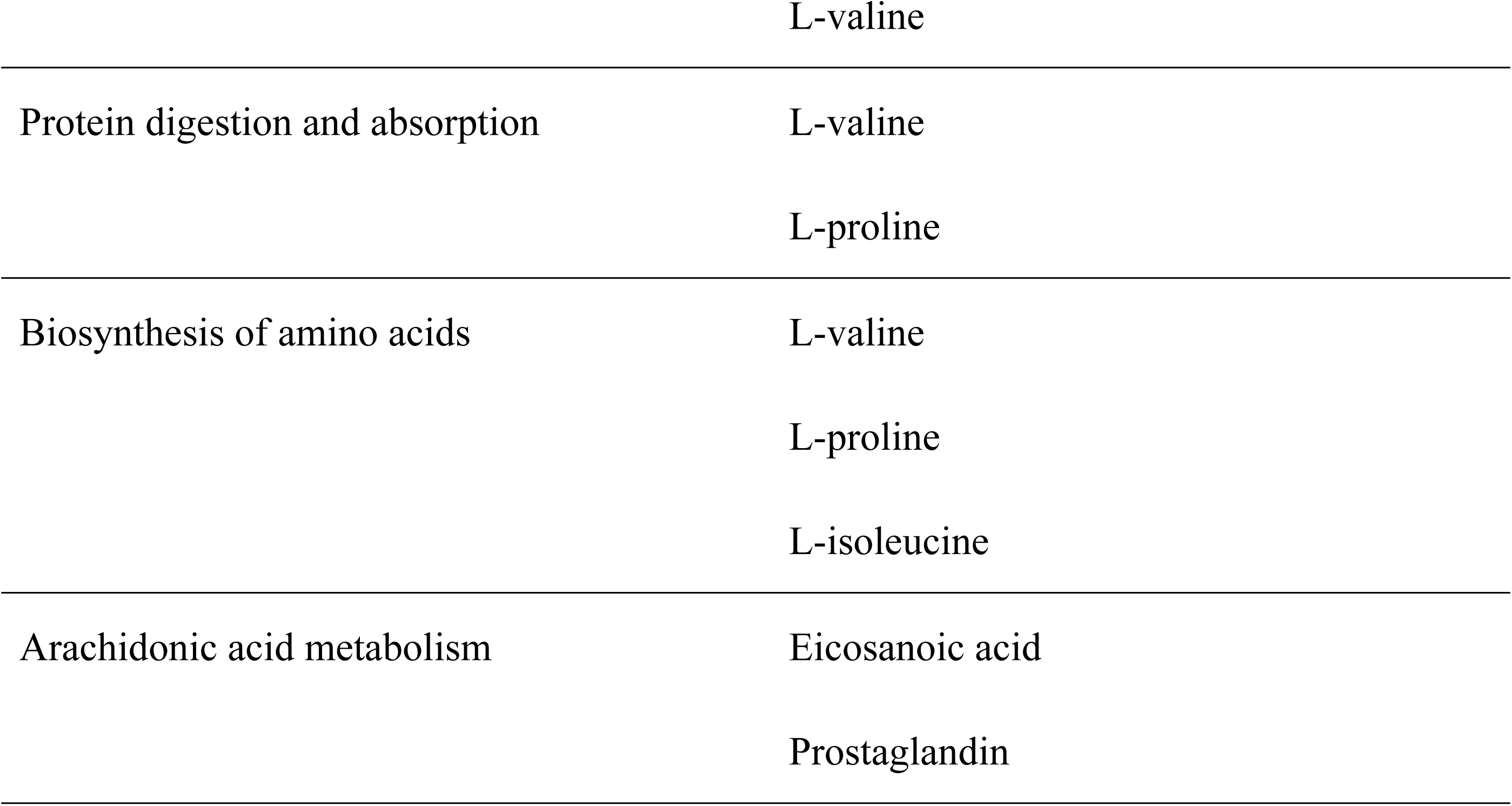
Kyoto Encyclopedia of Genes and Genomes pathway enrichment of different metabolites in skin tissues.

### Correlation analysis of genes and metabolites

To further determine the relationship between genes and metabolites, Pearson correlation coefficients were calculated based on both expression profiles of genes/metabolites and comprehensive review of literature related to villi (S3 Table). The results indicate a notable level of correlation between genes such as *KRT9*, *KRT36*, *KRT79*, *FGF18*, *AWAT2*, *MOGAT1*, *ACSM1*, *PM20D1*, and *SERPINE1* and metabolites such as arachidonic acid and melatonin; the Pearson correlation coefficients between the genes and arachidonic acid were all > 0.6.

## Discussion

As a barrier for organisms, the skin has a rich enzyme system and performs a range of transcriptional and metabolic activities, such as gene expression, oxidation, reduction, and hydrolysis [12]. Secondary organs of the skin, such as the sebaceous glands, sweat glands, and hair follicles, also possess specific transcription and metabolic capacity. The present study employed integrated transcriptomic and metabolomic analyses to elucidate the molecular mechanisms underlying cashmere quality in Liaoning cashmere goats. By comparing cashmere goats with coarse and fine hair, key DEGs and metabolites potentially involved in the regulation of SF development and cashmere fiber diameter were identified.

### Transcriptomic insights into skin tissue of cashmere goats

Transcriptomic analysis identified 211 DEGs between coarse and fine cashmere goats, with 75 genes upregulated and 136 genes downregulated in the coarse-haired group compared to in the fine-haired group. Notably, genes such as *FGF18*, *KRT36*, *KRT79*, and *AWAT2* were enriched significantly in pathways related to PPAR signaling and glycerol-lipid metabolism. The pathways play crucial roles in hair follicle development and sebum secretion, which are critical determinants of cashmere quality [13,14]. Upregulation of *FGF18* and *KRT9* in coarse- haired goats suggests their potential role in promoting thicker fiber growth.

*FGF18*, a member of the fibroblast growth factor family, regulates cell proliferation and differentiation in various tissues, including skin and hair follicles [15]. In cashmere goats, *FGF18* reportedly promotes the development and maintenance of hair follicles by stimulating the proliferation and differentiation of keratinocytes, the cells that produce keratin proteins [16]. In cashmere goats, the proper development of SFs is critical for high-quality cashmere production. *FGF18* upregulation in coarse-haired goats may enhance proliferation of keratinocytes and dermal papilla cells, leading to thicker fiber production.

Similarly, *KRT36* and *KRT79*, which encode keratin proteins, are crucial for the structural integrity of hair fibers [17], providing mechanical strength and resilience.

Downregulation of *KRT79* in coarse wool goats may lead to a decrease in the tensile strength of the fibers and a coarsening of the texture.

Two other notable DEGs in our study are *AWAT2* and *MOGAT1*, which are involved in lipid metabolism pathways. *AWAT2* encodes an enzyme involved in the synthesis of wax esters, which are essential components of sebum [18]. Downregulation of *AWAT2* in coarse cashmere goats implies that impaired lipid metabolism may negatively impact hair follicle health and fiber quality. The finding is supported by previous research demonstrating that lipid metabolism pathways, such as glycerol–lipid metabolism, are essential for the proper functioning of hair follicles and sebum secretion [19]. Similarly, *MOGAT1*, which is involved in triglyceride synthesis, may influence lipid metabolism in the skin, affecting the thickness and texture of cashmere fibers. The findings are consistent with the results of previous studies that have highlighted the importance of lipid metabolism in hair follicle development and fiber quality [4].

### Metabolomic study on skin tissue of cashmere goat

Metabolomic analysis identified 50 metabolites at significantly different levels in fine and coarse-haired goats, which were enriched mainly in ABC transporters, amino acid biosynthesis and metabolism pathways, arachidonic acid metabolism, protein digestion and absorption, and other pathways. Among them, L-proline and L-valine appear in multiple pathways.

Upregulation of amino acids such as proline and valine in coarse-haired goats suggests an enhanced protein synthesis capacity, which could contribute to thicker fiber production.

Proline is a key component of collagen, which is essential for the structural integrity of skin and hair follicles [20]. Upregulation of proline in coarse-haired goats may enhance collagen production, leading to stronger and thicker fibers. Similarly, valine, a branched-chain amino acid, is involved in protein synthesis and energy metabolism. Its upregulation in coarse-haired goats may support the energy demands of hair follicle growth and fiber production.

Arachidonic acid metabolism is another pathway that was significantly enriched in the present study. Arachidonic acid is a polyunsaturated fatty acid that plays a critical role in cellular signaling and inflammation regulation [21]. The involvement of this pathway in cashmere quality suggests that lipid-derived signaling molecules may influence hair follicle development and fiber properties. Specifically, arachidonic acid metabolites such as prostaglandins and leukotrienes reportedly modulate cellular functions, including proliferation and differentiation, which are essential for hair follicle growth [3]. The correlation between arachidonic acid metabolism and key genes identified in our study, such as *FGF18* and *KRT36*, further supports the notion that lipid signaling pathways are intricately linked to the structural components of hair fibers.

The enrichment of ABC transporter pathways highlights the importance of these transporters in regulating cellular ion channels and neuroendocrine functions. ABC transporters are involved in the transport of various molecules across cell membranes, including lipids, amino acids, and ions [3]. Upregulation of metabolites associated with ABC transporter pathways suggests that enhanced cellular transport mechanisms may contribute to the improved fiber quality observed in fine cashmere goats. The finding is consistent with previous studies demonstrating that ion channels and neuroendocrine signaling pathways indirectly regulate hair growth and development [3].

### Integration of Transcriptomics and Metabolomics

Integration of transcriptomics and metabolomics data in the present study provides a comprehensive view of the regulatory networks underlying cashmere quality. Our study identified significant correlations between key genes and metabolites, suggesting coordinated regulation of gene expression and metabolic pathways. The identification of key regulatory genes and metabolites provides valuable insights for developing targeted interventions to enhance cashmere production. This integrative approach has been applied successfully in previous studies to unravel complex biological processes [9,10]. Targeted breeding programs could focus on selecting goats with favorable genetic variants of genes such as *FGF18* and *KRT36* to enhance fiber fineness. Additionally, nutritional interventions aimed at modulating lipid metabolism and amino acid availability could be developed to optimize cashmere quality. For example, dietary supplementation with essential fatty acids or amino acids may promote the expression of key genes and metabolites identified in the present study, in turn improving cashmere yield and quality.

## Conclusion

In conclusion, the present study provides an analysis of the transcriptomic and metabolomic profiles associated with cashmere quality in Liaoning cashmere goats. The key regulatory genes and metabolites screened in the present study may provide insight into the molecular mechanisms underlying differences in cashmere fiber diameter. Future work should focus on validating the functional roles of the genes and metabolites through in vivo and in vitro experiments. Additionally, development of targeted breeding and nutritional strategies based on the findings could facilitate sustainable development of the cashmere goat industry.

## Data availability statement

The data that support the findings of this study are openly available in FigShare at http://doi.org/10.6084/m9.figshare.26886163 and http://doi.org/10.6084/m9.figshare.26886394.

## Acknowledgments

We would like to thank the Liaoning Modern Agricultural Production Base Construction Center at Liaoyang, China, for providing the test animals. We also thank Wei Ma and Chunqiang Wang for their valuable suggestions regarding this study. We would like to thank Editage (www.editage.cn) for English language editing.

## Supporting information captions

**S1 Table. Description of sequencing data.**

**S2 Table. Primer information for target gene amplification.**

**S3 Table. Pearson correlation coefficients indicating correlation between metabolomics and transcriptomics data (absolute value of correlation coefficient ≥ 0.6).**

**S1 File. Differentially Expressed Gene results.**

**S2 File. Gene Ontology enrichment results (Level 2).**

**S3 File. All differential metabolites.**

